# Of Microbes and Mange: Consistent changes in the skin microbiome of three canid species infected with sarcoptic mange

**DOI:** 10.1101/709436

**Authors:** Alexandra L. DeCandia, Kennedy N. Leverett, Bridgett M. vonHoldt

## Abstract

Sarcoptic mange is a highly contagious skin disease caused by the ectoparasitic mite, *Sarcoptes scabiei*. Although it afflicts over 100 mammal species worldwide, sarcoptic mange remains a disease obscured by variability at the individual, population, and species levels. Amid this variability, it is critical to identify consistent drivers of morbidity, particularly at the skin barrier. We characterized the skin microbiome of three species of North American canids: coyotes (*Canis latrans*), red foxes (*Vulpes vulpes*), and gray foxes (*Urocyon cinereoargenteus*). Comparing mange-infected and uninfected individuals, we found remarkably consistent signatures of microbial dysbiosis associated with mange infection. Across genera, mange-infected canids exhibited reduced microbial diversity, altered community composition, and increased abundance of opportunistic pathogens. The primary bacteria comprising these secondary infections were *Staphylococcus pseudintermedius*, previously associated with canid ear and skin infections, and *Corynebacterium spp*, previously found among the gut flora of *S. scabiei* mites and hematophagous arthropods. Considered together, this evidence suggests that mange infection consistently alters the canid skin microbiome and facilitates secondary bacterial infection. These results provide valuable insights into the pathogenesis of mange at the skin barrier of North American canids and can inspire novel treatment strategies. By further adopting a “One Health” framework that considers mites, microbes, and the potential for interspecies transmission, we can better elucidate the patterns and processes underlying this ubiquitous and enigmatic disease.

## INTRODUCTION

Sarcoptic mange has been termed a “ubiquitous neglected disease” (Hengge et al. 2006; Arlian & Morgan 2017). Although it afflicts over 100 mammal species on every continent except for Antarctica, numerous questions remain about its pathology (Arlian 1989; Bornstein et al. 2001; Walton et al. 2011; Astorga et al. 2018). A major impediment regards the wide-scale variability sarcoptic mange exhibits at every level of infection from individuals to populations to species, despite its universal source being *Sarctopes scabiei* mites (Pence & Ueckermann 2002).

Canids typify this variation. Considered prominent hosts of mange, many canid species are particularly susceptible due to their den usage and sociality (Bornstein et al. 2001; Kolodziej-Sobocinska et al. 2014; Montecino-Latorre et al. 2019). Yet individuals are not affected uniformly. Host symptoms range from mild pruritus to emaciation, dehydration, crust formation, or even death (Arlian 1989; Newman et al. 2002; Almberg et al. 2012; Nimmervoll et al. 2013). This variation scales to the population and species levels, where sarcoptic mange can exist as an enzootic parasite that imposes persistent, low levels of morbidity, or an epizootic parasite that causes dramatic mortality events in virulent outbreaks (Mörner 1992; Lindström et al. 1994; Pence & Windberg 1994; Gortázar et al. 1998; Little et al. 1998; Gosselink et al. 2007; Al-Sabi et al. 2014; Kolodziej-Sobocinska et al. 2014; Niedringhaus et al. 2019).

Amid this variability, it is important to elucidate consistent drivers of morbidity, particularly at the skin barrier. Considered the first line of defense against infection, the skin presents a physical and microbial barrier to invading parasites (Buffie & Pamer 2013; Grice 2014; Honda & Littman 2016). Upon contact with this barrier, adult females burrow into the skin to feed on host lymph and deposit the next generation of eggs (Hengge et al. 2006; Arlian & Morgan 2017). Often completing their entire life cycle on the same host, mites and their secretions continuously irritate the skin and elicit severe allergic reactions (Arlian 1989; Bornstein et al. 2001; Walton et al. 2004). Secondary bacterial infection with pathogenic microbes (such as *Staphylococcus spp*. and *Streptococcus spp*.) typically follows mite infestation (Walton et al. 2004; Walton 2010). Mites may even facilitate colonization of opportunistic invaders by transporting harmful bacteria to the host’s skin (Shelley et al. 1988) and enable proliferation by secreting immune inhibitors into burrows and lesions (Mika et al. 2012b; Swe & Fischer 2014).

To examine the effect of mite infection on the skin microbiome, Swe et al. (2014) experimentally infected pigs (*Sus scrofa domesticus*) with *S. scabiei* var. *suis* and sequenced microbial communities over the course of infection. Mange-infected individuals displayed lower levels of microbial diversity, altered community abundance, and increased incidence of *Staphylococcus spp*. compared to their uninfected counterparts. Similar patterns have been observed in humans, domestic animals, and wildlife infected with sarcoptic mange (Walton et al. 2004; Hengge et al. 2006; Almberg et al. 2012; Fraser et al. 2016), as well as domestic dogs (*Canis familiaris*) and humans with allergic skin conditions, such as atopic dermatitis (Kong et al. 2012; Rodrigues Hoffmann et al. 2014; Williams & Gallo 2015; Bradley et al. 2016; Wollina 2017). This evidence suggests that disrupted microbial communities may play a key role in the pathogenesis of sarcoptic mange.

Given the pervasive variability of this neglected disease, additional studies are needed to assess the universality of these trends. We contributed to these efforts by characterizing the skin microbiome of mange infection across three species of North American canids: coyotes (*Canis latrans*), red foxes (*Vulpes vulpes*), and gray foxes (*Urocyon cinereoargenteus*). Canids present an ideal system for these analyses, as they are among the primary species affected by sarcoptic mange in North America (Niedringhaus et al. 2019). Due to the divergent evolutionary histories of these three genera, we anticipated species-specific differences in microbial community composition of healthy and infected individuals. However, given their similar ecologies, we predicted consistent responses to mange infection that included decreased species richness and altered community abundance favoring pathogenic bacteria.

## MATERIALS & METHODS

### Sample and data collection

We opportunistically collected samples from coyotes, red foxes, and gray foxes admitted to licensed wildlife rehabilitation centers between January 2017 and April 2019. Partnering centers included the Wildlife Rehabilitation Center of Minnesota (MN), Fund for Animals Wildlife Center (CA), Janet L. Swanson Wildlife Health Center at Cornell University (NY), Woodlands Wildlife Refuge (NJ), PAWS Wildlife Center (WA), and Tufts Wildlife Clinic (MA). Critically, samples were collected upon admission to each facility and before treatment with antimicrobials, antivirals, anthelmintics, or acaricides. This minimized potential confounding effects of artificial environment (such as indoor facilities or human contact), sampling location, or treatment regime.

Sample metadata included sampling date and location, primary reason for admission, species, sex, age, weight, and mange status. We assessed mange severity by assigning each individual to a mange class corresponding to the percentage body area that exhibited symptoms, such as lesions, crusts, or alopecia. Uninfected individuals were assigned to Mange Class 0, with Mange Class 1 defined as 0-5% of the body covered, Mange Class 2 by 6-50%, and Mange Class 3 by more than 50%, following (Pence et al. 1983).

We collected swabs from five body sites (Figure 1) that included the external ear, dorsal flank, outer back leg, axilla, and groin. We used a sterile BBL™ swab to sample the skin at each body site, rotating the swab tip by 90° every 10 strokes for 40 swab strokes total (Rodrigues Hoffmann et al. 2014). Samples were stored at −80°C until DNA extraction. The Princeton University Institutional Animal Care and Use Committee reviewed and approved of all sample collection procedures (Princeton IACUC #2084A-16).

**Figure 1.**
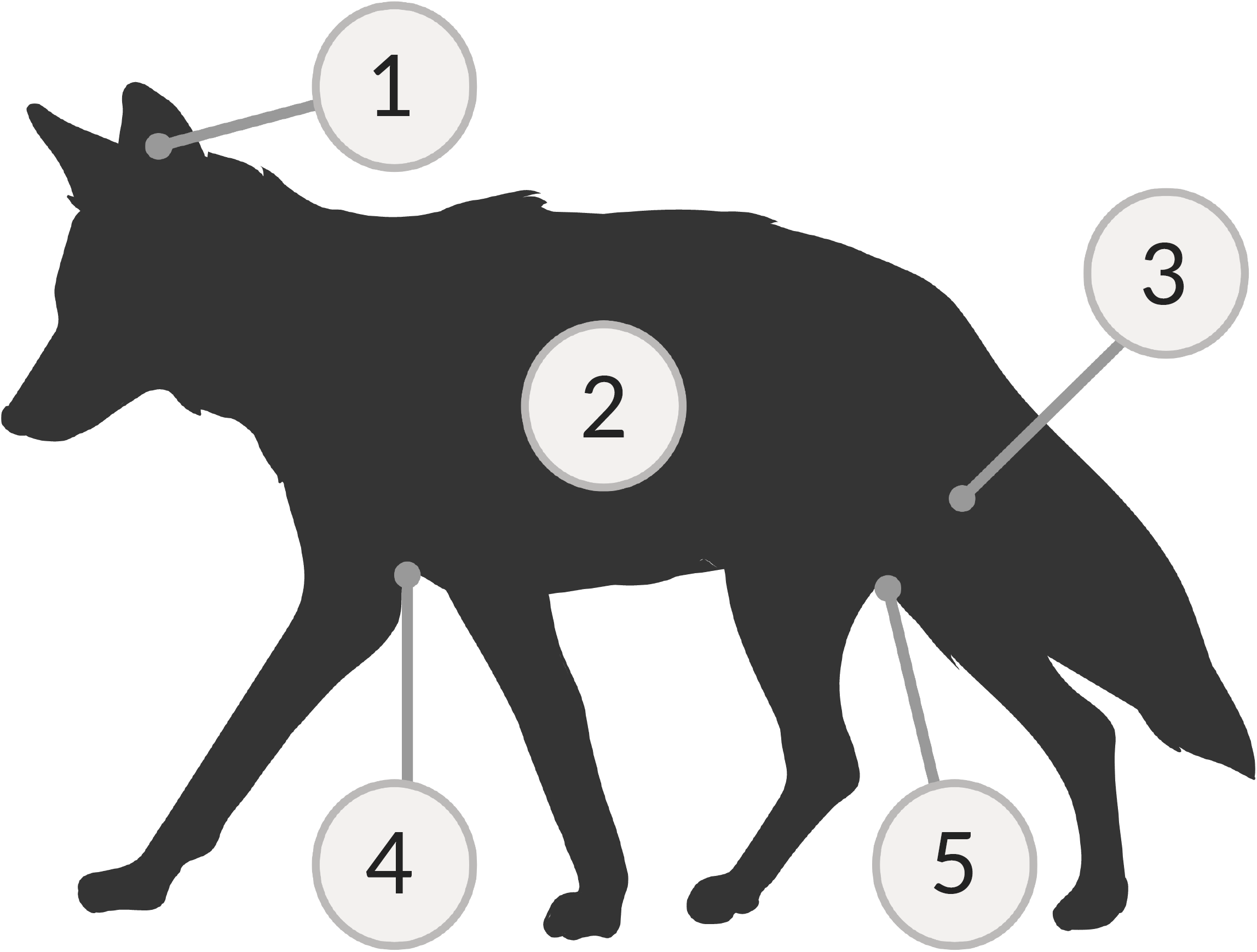
The five body sites swabbed included: (1) external ear, (2) dorsal flank, (3) outer back leg, (4) axilla, and (5) groin. Figure created with BioRender.

### DNA extraction and 16S rRNA V4 sequencing

We extracted microbial DNA from each swab tip using a modified DNeasy PowerSoil Kit (Qiagen Inc.) protocol described in (DeCandia et al. 2019). Briefly, we placed each swab tip into a PowerBead tube and used a Qiagen TissueLyser II to disrupt samples for two cycles, both 12 minutes at 20 shakes/second, with the addition of 60µL of C1 solution in between cycles. For the final elution step, we incubated samples at room temperature for 10-15 minutes using 60µL of C6 solution pre-heated to 70°C. We used sterile swab tips as negative controls during every round of extractions to minimize contamination risk. We subsequently concentrated extracts to 20µL in a Vacufuge and assessed DNA concentrations using a high-sensitivity Qubit™ fluorometer. We used molecular grade water to standardize samples to 2.5ng/µL and included low yield samples in subsequent steps.

We amplified and tagged the 16S ribosomal RNA (rRNA) hypervariable 4 (V4) region in each sample through polymerase chain reaction (PCR) using 96 unique combinations of barcoded forward (n=8) and reverse (n=12) primers (Caporaso et al. 2011). As in (DeCandia et al. 2019), the reaction recipe included 5µL HiFi HotStart ReadyMix (KAPA Biosystems), 3.2µL primer mix (1.25µM), and 1.8µL template DNA. Cycling conditions included: initial denaturation of 94°C/3 min, touchdown cycling for 30 cycles of [94°C/45s, 80-50°C/60s, 72°C/90s] decreasing 1°C each cycle, 12 cycles of [94°C/45s, 50°C/60s, 72°C/90s], and final extension of 72°C/10min. We used Quant-iT™ PicoGreen™ dsDNA assays to quantify PCR product, pooled equal nanograms of each library, and selected for amplicons between 300-400nt in length using Agencourt AMPure XP magnetic beads. We sent final libraries to the Princeton University Genomics Core Facility for paired-end amplicon sequencing (2×150nt) on an Illumina MiSeq.

### Data processing

We used a paired-end, dual-indexed barcode splitter implemented in *Galaxy* to demultiplex raw sequencing data, allowing for one nucleotide mismatch between expected and observed barcode sequences (Afgan et al. 2018). We then imported reads into *QIIME 2* v2019.4 (Caporaso et al. 2010; https://qiime2.org) for data filtering. Through the *dada2 denoise-paired* plugin, we corrected probable sequencing errors, removed chimeras, trimmed low quality bases, and merged paired-end reads to identify taxonomic features (Callahan et al. 2016). We additionally identified operational taxonomic units (OTUs) using *de-novo, closed reference*, and *open reference* clustering with *qiime vsearch* to compare our denoised dataset to more traditional cluster-based methods (Rognes et al. 2016).

### Alpha and beta diversity

We calculated alpha and beta diversity metrics using the *core-metrics-phylogenetic* and *alpha-rarefaction* functions in *QIIME 2*. To correct for differences in read depth, we rarefied samples to 5,153 sequences for the full dataset (n=125 samples) and 17,693 sequences for a composite dataset where samples were grouped by individual (n=25 grouped samples). Read depths were chosen to retain all samples for analysis.

To examine within-sample diversity, we calculated the Chao 1 Index for species richness and Pielou’s Evenness Metric for species equitability. For between-sample differences, we used *fasttree* to construct a rooted phylogenetic tree of taxonomic features and calculated Unweighted UniFrac distances for species presence, Weighted UniFrac distances for species presence and abundance, and the Bray-Curtis dissimilarity index for species abundance. We visualized sample dissimilarity through principal coordinate analysis (PCoA) using the *EMPeror* plugin (Vázquez-Baeza et al. 2013), and performed significance testing using the Kruskal-Wallis test for alpha diversity metrics and multivariate analysis of variance with permutation (PERMANOVA; Anderson 2001) for beta diversity differences. Variables of interest included sampling state, species, age, sex, year, and mange infection status.

### Taxonomic composition and differential abundance testing

We determined the taxonomic composition of each sample using a Naïve Bayes classifier trained on Greengenes 13_8 reference sequences trimmed to our 16S rRNA V4 amplicon and clustered at 99% similarity (DeSantis et al. 2006; Bokulich et al. 2018). We then used the *classify-sklearn* function to assign taxonomy to each representative sequence in the dataset (Bokulich et al. 2018).

To assess the statistical significance of compositional differences, we used two complementary approaches for differential abundance testing: analysis of composition of microbes (ANCOM) and gneiss balances. ANCOM calculates the log-ratio between pairwise combinations of taxa and sums how many times the null hypothesis is violated (Mandal et al. 2015). Gneiss calculates log transformed ratios (termed balances) between groups of taxa arranged in a hierarchical tree through correlation clustering (Morton et al. 2017). Ordinary least squares (OLS) regression can subsequently be used to test for differences between infection groups. Both analyses require a composition artifact as input, with additional filtering necessary to remove taxonomic features that occur in fewer than 10 samples or have frequencies below 50. We implemented each analysis with our composite dataset where samples were grouped by individual, and queried results using the NCBI BLASTn online tool (Altschul et al. 1990).

## RESULTS

### Amplicon sequencing and data processing

We sequenced 153 samples collected from 15 coyotes (mange-infected=9, uninfected=5, unknown=1), 13 red foxes (mange-infected=8, uninfected=5), and 2 gray foxes (mange-infected=1, uninfected=1). The full dataset contained 4,397,629 raw reads, which reduced to 3,911,712 sequences after denoising (Supplemental Table S1). The denoised dataset contained 11,800 unique taxonomic features, whereas the OTU datasets contained 6,137 (*de-novo)*, 5,456 (*closed reference)*, and 8,106 (*open reference*) features at 97% percentage identity. Proceeding with the denoised dataset, we removed 28 samples due to incorrect body sites (n=7), treatment prior to sampling (n=11), low read counts (n=5), and unknown mange status (n=5). Our final dataset consisted of 125 samples collected from 12 coyotes (mange-infected=8, uninfected=4), 11 red foxes (mange-infected=6, uninfected=5), and 2 gray foxes (mange-infected=1, uninfected=1).

### Uninfected samples cluster by individual rather than body site

Given repeated measures across individuals (n=5 samples per individual) and body sites (n=25 samples per body site) in the denoised dataset, we implemented principal coordinate analysis (PCoA) on uninfected samples to assess whether these factors significantly influenced beta diversity. Across all three distance measures, samples clustered by individual (PERMANOVA; Bray-Curtis, *pseudo-F*= 2.984, *p*=0.001; Unweighted UniFrac, *pseudo-F-* =2.938, *p*=0.001; Weighted UniFrac, *pseudo-F*=3.470, *p*=0.001) rather than body site (Bray-Curtis, *pseudo-F*=0.781, *p*=0.997; Unweighted UniFrac, *pseudo-F*= 0.769, *p*=0.997; Weighted UniFrac, *pseudo-F*=0.950, *p*=0.574; Figure 2, Supplemental Figure S1). We therefore grouped samples by individual in downstream analyses to control for statistical relicts of pseudoreplication. Rather than five samples per canid (*i*.*e*., one for each body site), each individual was represented by one composite sample that contained all features in their skin microbiome.

**Figure 2.**
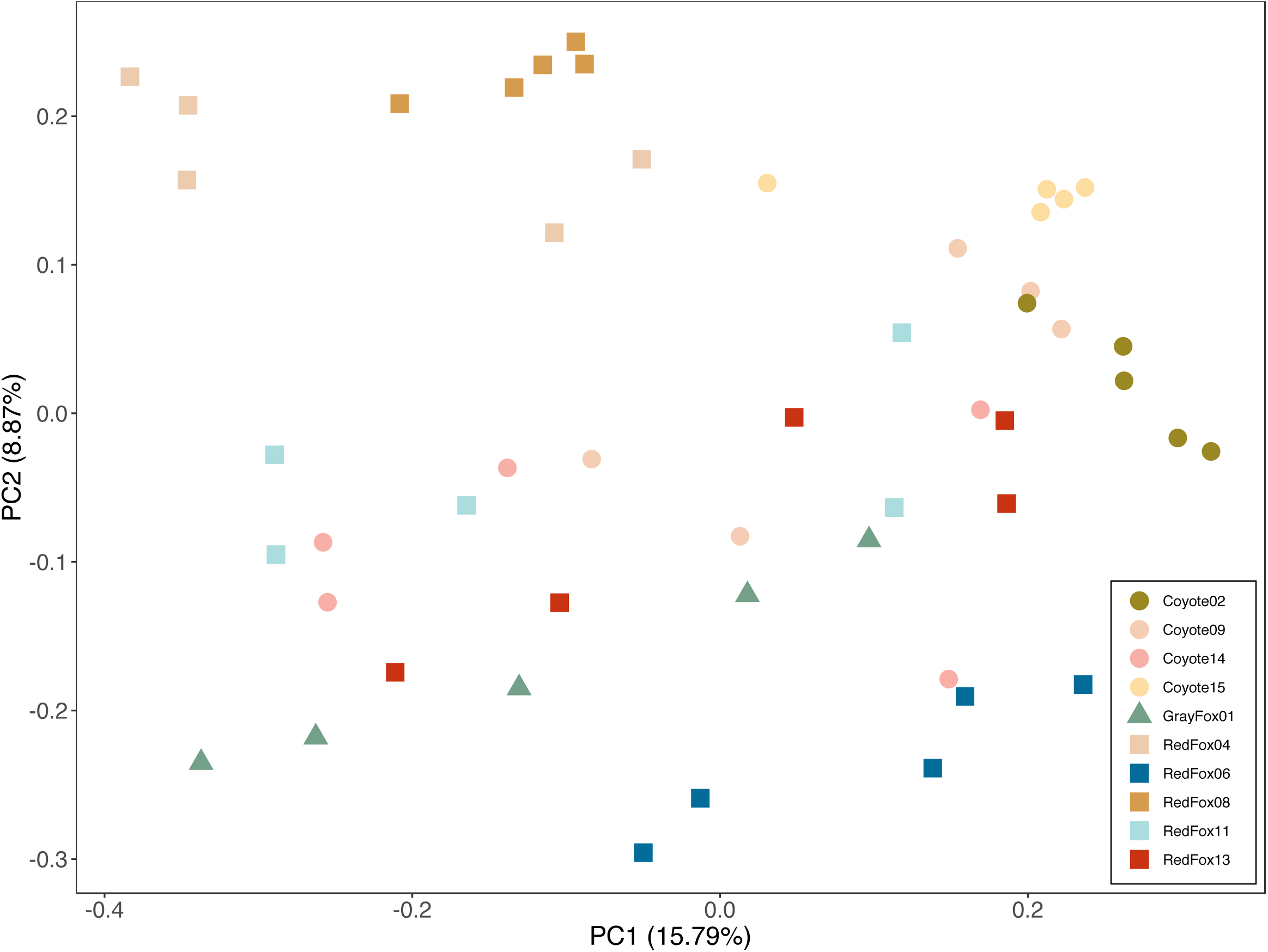
Principal coordinate analysis (PCoA) of uninfected individuals showed significant clustering by individual (PERMANOVA; *pseudo-F*=2.938, *p*=0.001) rather than body site (*pseudo-F*= 0.769, *p*=0.997) using phylogeny-based Unweighted UniFrac distances.

We performed significance testing for alpha and beta diversity on our composite dataset to determine which metadata categories were predictive of microbial community structure. Across metrics, mange infection was the variable most strongly associated with differences in alpha and beta diversity (Supplemental Table S2). Although sex appeared significant, further analyses showed non-independence between sex and mange status (Chi-square test, *X*^*2*^=4.039, *p*=0.04), due to a disproportionate number of infected males. All other categories exerted minimal effects of community structure. We therefore analyzed the full composite dataset for subsequent analyses and used mange infection status as our variable of interest.

### Mange-infected canids exhibit decreased diversity and community evenness across species

We observed significantly reduced species richness (Kruskal-Wallis test; Chao 1 Index, *H*=10.711, *p*=0.001; Figure 3A) and evenness (Pielou’s Eveness Metric, *H*= 8.643, *p*=0.003; Figure 3B) in mange-infected individuals. Beta diversity similarly differed by infection group. Measures of species abundance (PERMANOVA; Bray-Curtis, *pseudo-F*=3.885, *p*=0.001; Figure 3C), presence (Unweighted UniFrac, *pseudo-F*=2.211, *p*=0.006; Supplemental Figure S2A), and both presence and abundance considered together (Weighted UniFrac, *pseudo-F*=4.398, *p*=0.001; Supplemental Figure S2B) showed significant differences between mange-infected and uninfected canids. For all three measures, samples clustered by infection status along PC1, which explained 16.49-29.01% of the variation.

**Figure 3.**
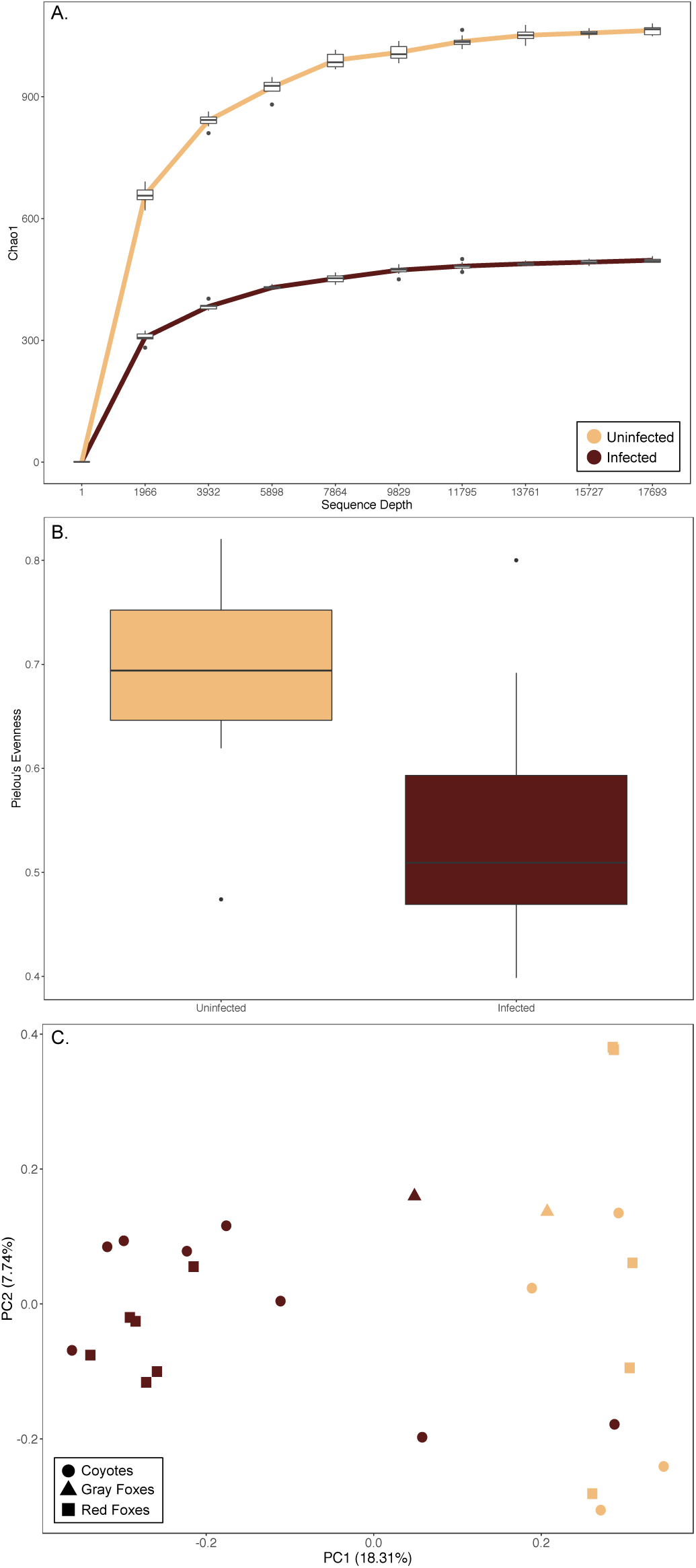
Mange-infected individuals had significantly reduced (A) species richness (Kruskal-Wallis test; Chao 1, *H*=10.711, *p*=0.001) and (B) eveness (Pielou’s Evenness Metric, *H*=8.643, *p*=0.003) when compared to uninfected individuals. (C) Beta diversity also differed significantly between infection groups (PERMANOVA; Bray-Curtis, *pseudo-F*=3.885, *p*=0.001).

Taxonomic composition of skin microbial communities confirmed these patterns (Figure 4). Although variation between individuals was evident, mange-infected canids exhibited higher relative abundance of Actinobacteria (infected=25.848%, uninfected=11.843%) and Bacilli (infected=35.839%, uninfected=9.902%), and reduced abundance of “Other” taxa (infected=7.520%, uninfected=25.828%). These results remained consistent even when the dataset was subdivided by species (Supplemental Table S3).

**Figure 4.**
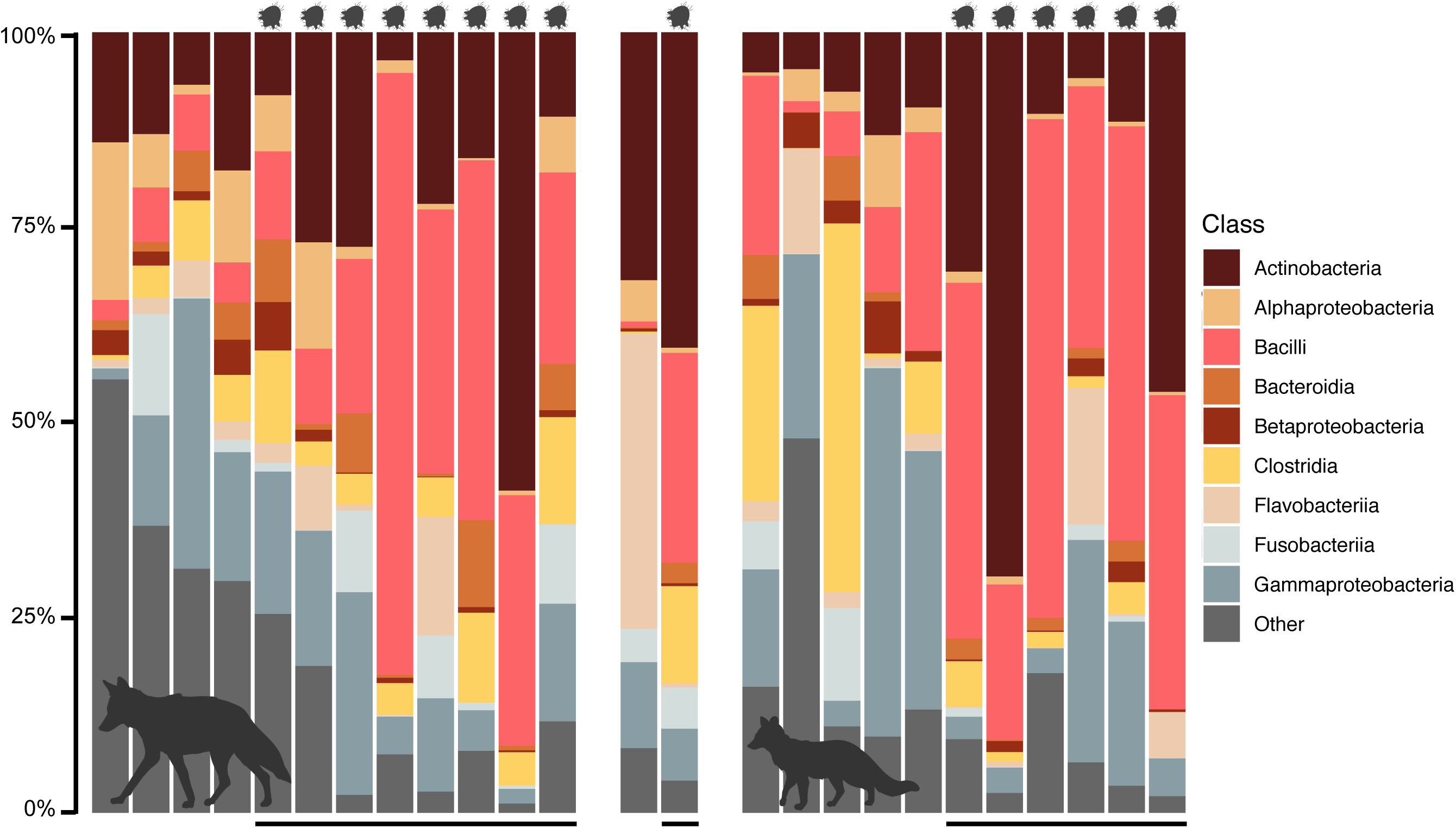
Taxonomic composition of skin microbial communities for 12 coyotes, 2 gray foxes, and 11 red foxes. Black bars (bottom) and mites (top) signify individuals infected with sarcoptic mange. Figure created with BioRender.

### Increased abundance of Staphylococcus pseudintermedius and Corynebacterium spp. with mange infection

Analysis of the composition of microbes (ANCOM) returned one taxonomic feature as consistently and significantly more abundant in mange-infected individuals: feature 3f0449c545626dd14b585e9c7b2d16f4 (*W*=111; Supplemental Figure S3). NCBI BLASTn (Altschul et al. 1990) search results returned high sequence similarity to *Staphylococcus pseudintermedius* (class: Bacilli; Supplemental Table S4A). Although not statistically significant, feature e3e89166daa575e51d7a14bc65f11153 exhibited the second highest number of rejected null hypotheses (*W*=21) and matched *Corynebacterium spp*. (class: Actinobacteria; Supplemental Table S4B).

Given the strong effect of mange infection on alpha and beta diversity, we constructed a simple OLS regression model using mange infection and gneiss balances as our variables of interest. This model explained 9.40% of the variation observed, and returned two statistically significant balances that exhibited increased taxonomic abundance in mange-infected individuals: y02 and y05 (both *p*=0.013). After visualizing the tree hierarchy through the Interactive Tree of Life (iTOL) v3 online tool (Letunic & Bork 2016), we found that balance y05 was nested within y02. As a result, both balances pointed towards the same signal: increased proportion of features 3f0449c545626dd14b585e9c7b2d16f4 (mean infected=0.421, uninfected=0.032) and e3e89166daa575e51d7a14bc65f11153 (mean infected=0.170, uninfected=0.003) in mange-infected individuals (Figure 5A). These features were previously identified as *S. pseudintermedius* and *Corynebacterium spp*. using NCBI BLASTn, and were clustered with two additional features in the dendrogram relating all taxa: features c2d41dc0a7b8eaedcf4697512aee4427 (identified as *Staphylococcus spp*.) and 22a5bce17370d6c495f5e83232650ec7 (identified as *Streptococcus agalactiae*; Figure 5B). These additional features exhibited higher proportions in mange-infected canids (*Staphylococcus spp*. mean infected=0.017, uninfected=0.001; *S. agalactiae* mean infected=0.007, uninfected<0.001). Although balance y78 was also statistically significant, its proportions only marginally differed between infection groups, with increased abundance of its component taxa found in uninfected canids.

**Figure 5.**
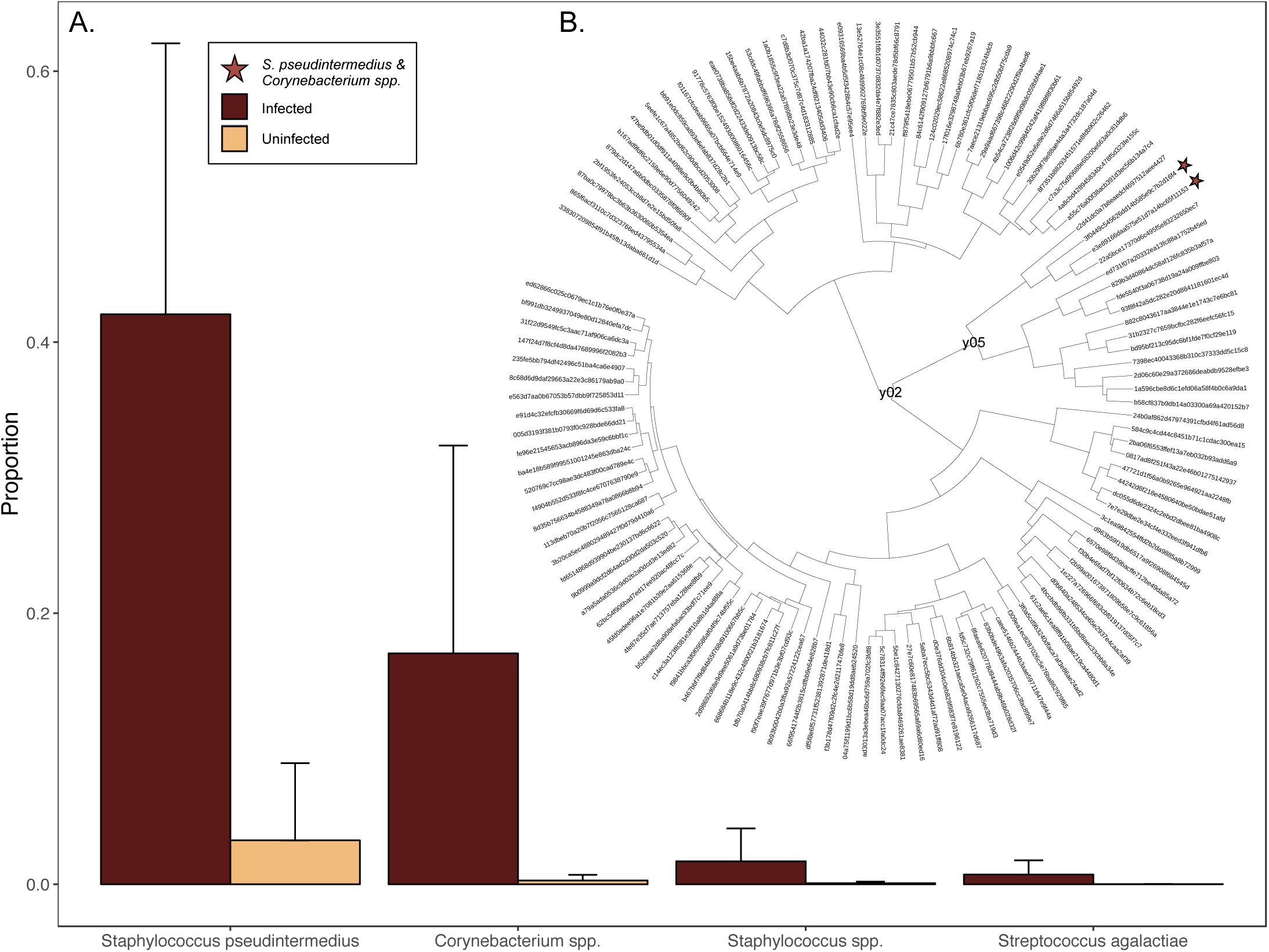
(A) Relative abundance of four taxonomic features found within gneiss balances associated with sarcoptic mange infection. *Staphylococcus pseudintermedius* and *Corynebacterium spp*. exhibited the largest differences between infection groups, with *Staphylococcus spp*. and *Streptococcus agalactiae* clustered with these taxa in the (B) hierarchy relating all features through correlation clustering.

## DISCUSSION

Sarcoptic mange is among the most widespread diseases affecting mammals on a global scale. Despite recognition since antiquity (Arlian & Morgan 2017), mange is considered a neglected disease, as there remain numerous questions about its pathology in free ranging wildlife (Astorga et al. 2018). The interplay between mites and microbes at the skin barrier is one such question, given increasing recognition of the importance of host-associated microbiomes in wildlife health and disease (DeCandia et al. 2018; Hauffe & Barelli 2019; Trevelline et al. 2019).

We characterized the skin microbiome of mange-infected and uninfected canids in three North American species: coyotes, red foxes, and gray foxes. Across species, we observed remarkably consistent signatures of mange infection that included reduced diversity, shifted community composition, and increased incidence of *S. pseudintermedius* and *Corynebacterium spp*. Although samples derived from different species sampled in different states, infection status was the primary driver of microbial community structure in terms of species richness, evenness, presence, and relative abundance.

Commensal microbial communities are shaped by a complex milieu of genetic and environmental factors (Bonder et al. 2016; Rothschild et al. 2018). Although inter-individual variation is pervasive, the host-associated microbiome is thought to exhibit phylosymbiosis between microbes and their hosts over evolutionary timescales (Brucker & Bordenstein 2012; Brooks et al. 2017). In a study of small mammals spanning six genera, for example, species identity exerted a far stronger effect on microbial community structure than did local habitat (Knowles et al. 2019). We therefore anticipated divergence between the skin microbiome of our three focal species, as coyotes, red foxes, and gray foxes derive from different genera within Canidae.

Counter to this expectation, we found minimal differences between skin microbial communities across species, sampling locations, years, and ages. Instead, mange infection status was the primary factor associated with microbial community structure within our dataset. This suggested two primary hypotheses. The first posits that shared evolutionary history and contemporary ecology of these species leads to similar skin microbiomes, as seen in gut microbial communities across families within class Mammalia (Nishida & Ochman 2018). The second contends that mange infection alters community composition consistently and dramatically across species, thereby blurring inter-genus distinctions within our relatively small sample set.

Results from this study primarily supported the second hypothesis, although it is likely that evolutionary history, contemporary ecology, and mange infection all influence the observed patterns of microbial diversity. Within the broader context of microbes and mange, reduced microbial variation and increased abundance of opportunistic pathogens is consistent with humans infected with *S. scabiei* var. *hominis* (McCarthy et al. 2004; Whitehall et al. 2013), pigs experimentally infected with *S. scabiei* var. *suis* (Swe et al. 2014), Santa Catalina island foxes (*Urocyon littoralis catalinae*) infected with *Otodectes cynotis* ear mites (DeCandia et al. 2019), and domestic dogs and humans exhibiting allergic skin disorders (Kong et al. 2012; Hoffmann et al. 2014; Williams & Gallo 2015; Bradley et al. 2016; Wollina 2017). Although the identity of opportunistic pathogens varies by host species, *Staphylococcus spp*. and *Streptococcus spp*. were commonly reported. Mite presence may even facilitate these secondary bacterial infections by secreting proteins that inhibit the mammalian complement system, known to be a key player in the immune response against mite and bacterial infections (Bergström et al. 2009; Mika et al. 2012b, 2012a; Swe & Fischer 2014). Mite burrows and host lesions may therefore provide ideal environments for opportunistic pathogens to proliferate.

The primary microbial taxa associated with mange infection in this study included *S. pseudintermedius* and *Corynebacterium spp*., with *S. agalactiae* and other *Staphylococcus spp*. marginally differing in abundance. Both humans and pigs infected with *S. scabiei* exhibited increased proportion of *S. aureus* (Whitehall et al. 2013; Swe et al. 2014), with *S. pseudintermedius* reported in island foxes infected with ear mites (DeCandia et al. 2019). These analogous results present compelling evidence that mite infection is associated with *Staphylococcus spp*. proliferation across host taxa. Further, increased abundance of *S. pseudintermedius* across four canid species infected with *S. scabiei* (coyotes, red foxes, and gray foxes) and *O. cynotis* (island foxes; DeCandia et al. 2019) mites suggests that it is an important bacterial taxon within Canidae.

A common canid commensal (Bannoehr et al. 2009), *S. pseudintermedius* becomes an opportunistic pathogen when the skin microbiome is disrupted by allergic skin disease, infection, or surgery (Fazakerley et al. 2009; Bannoehr & Guardabassi 2012; Ngo et al. 2018). Resultant biofilms lead to chronic inflammation in domestic dogs, cats (*Felis catus*), and to a lesser extent, humans (Pompilio et al. 2015), with antibiotic resistant strains emerging across veterinary and medical hospitals (Sasaki et al. 2007; Weese & van Duijkeren 2010).

Although less commonly reported across host species, *Corynebacterium spp*. was detected in skin crusts and *S. scabiei* mites isolated from pigs with severe mange (Swe et al. 2014). Similar bacteria were also isolated from the gastrointestinal tracts of hematophagous arthropods, such as triatomes (*Triatoma infestans*; Durvasula et al. 2008) and three species of ticks (*Ixodes ricinus, Dermacentor reticulatus*, and *Haemaphysalis concinna*; Rudolf et al. 2009). This evidence suggests that *Corynebacterium spp*. may derive from mite bodies, secretions, or frass deposited at the site of infection, in addition to canid commensal communities. As with *S. pseudintermedius*, these bacteria likely benefit from mite inhibition of mammalian complement.

These additional insights into the pathogenesis of sarcoptic mange may enable novel management of infected wildlife *in* and *ex situ* (Rowe et al. 2019). Regarding treatment, commonly used acaricides possess harmful side effects for individuals and the environment, while drug resistance has been observed in *S. scabiei* mites and their concomitant bacterial infections (Walton et al. 2004; Hengge et al. 2006; Weese & van Duijkeren 2010; Thomas et al. 2015). It may become critical to pursue novel avenues of treatment, such as anti- or probiotic therapies, to ensure long-term control of this disease. Insights into mite microbiomes may further provide means of mite control if these communities can be manipulated (Durvasula et al. 2008).

Given the ubiquity of this disease and its capacity to infect humans, domestic animals, and wildlife, mange presents an ideal candidate for adopting a “One Health” perspective when mitigating its negative effects (Astorga et al. 2018). Mammalian hosts can be intricately coupled, enabling interspecies transmission when diseased animals approach human settlements in search of resources or shelter, as seen in mange-infected coyotes (Murray et al. 2015; Murray & St. Clair 2017) and red foxes (Carricondo-Sanchez et al. 2017). Although public health concerns are minor due to the lesser severity of zoonotic mange, interspecies transmission between widespread and at risk species can pose a conservation risk. Thus identifying consistent drivers of morbidity, such as altered microbial communities, can enable better prediction and mitigation of mange dynamics across host systems.

## ACKNOWLEDGEMENTS

We would like to thank Leslie Reed from the Wildlife Rehabilitation Center of Minnesota (MN), Gina Taylor and Ali Crumpacker from the Fund for Animals Wildlife Center (CA), Sara Childs-Sanford from the Janet L. Swanson Wildlife Health Center at Cornell University (NY), Heather Freeman from the Woodlands Wildlife Refuge (NJ), Emily Meredith from PAWS Wildlife Center (WA), and Florina Tseng from the Tufts Wildlife Clinic (MA) for contributing samples. We would additionally like to thank Lindy McBride, Wei Wang, Jessica Wiggins, and Edward Schrom from Princeton University for their support during library preparation, sequencing, and data processing. Funding for this study was provided by the American Society of Mammalogists Grants-in-Aid of Research program. This material is based upon work supported by the National Science Foundation Graduate Research Fellowship under Grant No. DGE1656466.

## DATA ACCESSIBILITY

Upon acceptance, demultiplexed reads will be deposited to the NCBI Sequence Read Archive and sample metadata will be made available through supplementary materials.

## AUTHOR CONTRIBUTIONS

ALD and BMvH designed the study; ALD and KNL conducted the laboratory work; ALD processed and analyzed the data; ALD prepared the manuscript; ALD, KNL, and BMvH contributed to and approved the final manuscript.

