## Supplemental for "Of Microbes and Mange: Consistent changes in the skin microbiome of three canid species infected with sarcoptic mange"

### SUPPLEMENTAL TABLES & FIGURES

**Supplemental Table S1.** Sample metadata and sequence processing statistics for each stage of the *dada2* denoising pipeline implemented in *QIIME2* v2019.4.

| Sample | Individual | Body Site | Mange | Raw | Filtered | Denoised | Merged | Non-chimeric |
|---|---|---|---|---|---|---|---|---|
| Sample001 | Coyote01 | Axilla | N | 21880 | 21460 | 21211 | 20323 | 20318 |
| Sample002 | Coyote01 | Flank | N | 19921 | 19224 | 18726 | 17056 | 17053 |
| Sample003 | Coyote01 | Ext. Ear | N | 23282 | 22705 | 22189 | 20714 | 20702 |
| Sample004 | Coyote01 | Groin | N | 22307 | 21460 | 20996 | 18645 | 18597 |
| Sample005 | Coyote01 | Leg | N | 14010 | 13740 | 13284 | 12113 | 12110 |
| Sample006 | Coyote01 | Feces | N | 42790 | 41668 | 41450 | 40176 | 39773 |
| Sample007 | Coyote02 | Axilla | N | 34661 | 33742 | 32714 | 28442 | 28374 |
| Sample008 | Coyote02 | Flank | N | 32241 | 31450 | 30596 | 25559 | 25474 |
| Sample009 | Coyote02 | Ext. Ear | N | 26067 | 25433 | 24612 | 20560 | 20421 |
| Sample010 | Coyote02 | Groin | N | 35941 | 34529 | 33286 | 27620 | 27589 |
| Sample011 | Coyote02 | Leg | N | 24420 | 23845 | 23372 | 20910 | 20910 |
| Sample012 | Coyote03 | Axilla | Y | 28306 | 27017 | 25999 | 21460 | 21384 |
| Sample013 | Coyote03 | Flank | Y | 31527 | 30782 | 29924 | 26527 | 26508 |
| Sample014 | Coyote03 | Ext. Ear | Y | 27237 | 26792 | 25758 | 22549 | 22542 |
| Sample015 | Coyote03 | Groin | Y | 33222 | 32677 | 31576 | 27342 | 27269 |
| Sample016 | Coyote03 | Leg | Y | 27294 | 26883 | 26071 | 23309 | 23284 |
| Sample017 | Coyote04 | Axilla | Y | 23290 | 22464 | 21870 | 19271 | 19224 |
| Sample018 | Coyote04 | Flank | Y | 24863 | 24429 | 23722 | 21659 | 21584 |
| Sample019 | Coyote04 | Ext. Ear | Y | 22292 | 21937 | 21339 | 19382 | 19306 |
| Sample020 | Coyote04 | Groin | Y | 26914 | 26394 | 25780 | 24032 | 23980 |
| Sample021 | Coyote04 | Leg | Y | 35549 | 35049 | 34300 | 31853 | 31753 |
| Sample022 | Coyote05 | Axilla | Unk | 33548 | 32422 | 31613 | 28500 | 28491 |
| Sample023 | Coyote05 | Flank | Unk | 38426 | 37151 | 36068 | 32869 | 32798 |
| Sample024 | Coyote05 | Ext. Ear | Unk | 28579 | 27554 | 26875 | 24312 | 24258 |
| Sample025 | Coyote05 | Groin | Unk | 46654 | 45061 | 43252 | 36312 | 36189 |
| Sample026 | Coyote05 | Leg | Unk | 26783 | 25892 | 25109 | 22394 | 22219 |
| Sample027 | Coyote06 | Axilla | Y | 30134 | 29327 | 29144 | 28481 | 28119 |

| | | | | | | | | |
|---|---|---|---|---|---|---|---|---|
| Sample028 | Coyote06 | Flank | Y | 29076 | 28468 | 28300 | 27678 | 27559 |
| Sample029 | Coyote06 | Ext. Ear | Y | 48530 | 47202 | 46647 | 43752 | 43505 |
| Sample030 | Coyote06 | Groin | Y | 21978 | 21428 | 21220 | 20313 | 19973 |
| Sample031 | Coyote06 | Leg | Y | 33429 | 32852 | 32661 | 31772 | 31703 |
| Sample032 | Coyote07 | Axilla | Y | 37741 | 37029 | 36899 | 35555 | 34737 |
| Sample033 | Coyote07 | Flank | Y | 40204 | 39358 | 38854 | 36406 | 35824 |
| Sample034 | Coyote07 | Ext. Ear | Y | 27471 | 26564 | 25704 | 22593 | 22509 |
| Sample035 | Coyote07 | Groin | Y | 29636 | 28091 | 27972 | 25960 | 24267 |
| Sample036 | Coyote07 | Leg | Y | 14629 | 14270 | 13999 | 13233 | 13233 |
| Sample037 | Coyote08 | Axilla | Y | 43555 | 42887 | 42708 | 41674 | 40514 |
| Sample038 | Coyote08 | Flank | Y | 23519 | 23140 | 23053 | 22648 | 22328 |
| Sample039 | Coyote08 | Ext. Ear | Y | 62047 | 60594 | 60202 | 58765 | 58646 |
| Sample040 | Coyote08 | Groin | Y | 27548 | 26938 | 26803 | 26050 | 25592 |
| Sample041 | Coyote08 | Leg | Y | 40075 | 38521 | 37911 | 35059 | 34588 |
| Sample042 | Coyote09 | Axilla | N | 38705 | 38124 | 37924 | 37177 | 37040 |
| Sample043 | Coyote09 | Flank | N | 29913 | 29290 | 28450 | 25421 | 25414 |
| Sample044 | Coyote09 | Ext. Ear | N | 32290 | 31807 | 31315 | 30053 | 29800 |
| Sample045 | Coyote09 | Groin | N | 38723 | 37498 | 36734 | 33413 | 33402 |
| Sample046 | Coyote09 | Leg | N | 31017 | 30329 | 29494 | 25584 | 25584 |
| Sample047 | Coyote10 | Axilla | Y | 24923 | 24459 | 24276 | 23821 | 23665 |
| Sample048 | Coyote10 | Flank | Y | 34535 | 33325 | 33010 | 32030 | 31969 |
| Sample049 | Coyote10 | Ext. Ear | Y | 25538 | 25053 | 24828 | 24091 | 24036 |
| Sample050 | Coyote10 | Groin | Y | 26185 | 25455 | 25295 | 24581 | 24542 |
| Sample051 | Coyote10 | Leg | Y | 23103 | 22689 | 22610 | 22289 | 22289 |
| Sample052 | Coyote11 | Axilla | Y | 26320 | 25733 | 25586 | 24429 | 24208 |
| Sample053 | Coyote11 | Flank | Y | 26473 | 25880 | 25638 | 24280 | 23976 |
| Sample054 | Coyote11 | Ext. Ear | Y | 24999 | 24379 | 24093 | 22787 | 22536 |
| Sample055 | Coyote11 | Groin | Y | 47912 | 46327 | 46058 | 44134 | 42606 |
| Sample056 | Coyote11 | Leg | Y | 27332 | 26843 | 26535 | 25767 | 25247 |
| Sample057 | Coyote12 | Axilla | Y | 22901 | 22411 | 22067 | 20689 | 20689 |
| Sample058 | Coyote12 | Flank | Y | 9160 | 8849 | 8676 | 8146 | 8112 |
| Sample059 | Coyote12 | Ext. Ear | Y | 37803 | 36633 | 35937 | 34136 | 34032 |
| Sample060 | Coyote12 | Groin | Y | 17988 | 17586 | 17046 | 16172 | 16135 |

| | | | | | | | | |
|---|---|---|---|---|---|---|---|---|
| Sample061 | Coyote12 | Leg | Y | 21884 | 21394 | 20977 | 19999 | 19997 |
| Sample062 | Coyote13 | Axilla | Y | 318 | 276 | 216 | 152 | 152 |
| Sample063 | Coyote13 | Flank | Y | 24762 | 24139 | 23825 | 22783 | 22187 |
| Sample064 | Coyote13 | Ext. Ear | Y | 24572 | 23710 | 23344 | 20675 | 20580 |
| Sample065 | Coyote13 | Groin | Y | 44941 | 43545 | 43390 | 42686 | 41686 |
| Sample066 | Coyote13 | Leg | Y | 27261 | 26172 | 25972 | 25207 | 25038 |
| Sample067 | Coyote14 | Axilla | N | 24867 | 24344 | 24165 | 23564 | 23365 |
| Sample068 | Coyote14 | Flank | N | 34128 | 33267 | 32865 | 31360 | 31233 |
| Sample069 | Coyote14 | Ext. Ear | N | 23589 | 22874 | 22206 | 19363 | 19348 |
| Sample070 | Coyote14 | Groin | N | 20079 | 19648 | 19171 | 17101 | 17087 |
| Sample071 | Coyote14 | Leg | N | 27085 | 25835 | 25475 | 24560 | 24541 |
| Sample072 | Coyote15 | Axilla | N | 24297 | 23952 | 22981 | 19613 | 19541 |
| Sample073 | Coyote15 | Flank | N | 36185 | 35588 | 34415 | 30440 | 30248 |
| Sample074 | Coyote15 | Ext. Ear | N | 27551 | 27153 | 26573 | 24617 | 24431 |
| Sample075 | Coyote15 | Groin | N | 21389 | 20999 | 19923 | 16392 | 16313 |
| Sample076 | Coyote15 | Leg | N | 34751 | 33306 | 31768 | 24774 | 24475 |
| Sample077 | GrayFox01 | Axilla | N | 26003 | 25431 | 25015 | 22566 | 22542 |
| Sample078 | GrayFox01 | Flank | N | 36943 | 36220 | 36026 | 35639 | 35448 |
| Sample079 | GrayFox01 | Ext. Ear | N | 10981 | 10579 | 10370 | 9937 | 9937 |
| Sample080 | GrayFox01 | Groin | N | 6211 | 6048 | 5863 | 5289 | 5289 |
| Sample081 | GrayFox01 | Leg | N | 37515 | 37011 | 36839 | 36508 | 36249 |
| Sample082 | GrayFox02 | Axilla | Y | 24511 | 23550 | 23393 | 21973 | 21537 |
| Sample083 | GrayFox02 | Flank | Y | 18440 | 18094 | 17959 | 17572 | 17391 |
| Sample084 | GrayFox02 | Ext. Ear | Y | 22054 | 21469 | 21267 | 20437 | 20335 |
| Sample085 | GrayFox02 | Groin | Y | 36630 | 35789 | 35618 | 34348 | 33341 |
| Sample086 | GrayFox02 | Leg | Y | 30891 | 29825 | 29616 | 28345 | 28075 |
| Sample087 | RedFox01 | Center Back | Y | 10707 | 10351 | 10068 | 9525 | 9522 |
| Sample088 | RedFox01 | Ear | Y | 8401 | 8176 | 7961 | 7363 | 7360 |
| Sample089 | RedFox01 | Perianal | Y | 11617 | 11276 | 11170 | 10844 | 10844 |
| Sample090 | RedFox01 | Lip | Y | 16955 | 16613 | 16482 | 16155 | 16116 |
| Sample091 | RedFox01 | Nose | Y | 7390 | 7097 | 6952 | 6451 | 6440 |
| Sample092 | RedFox02 | Axilla | Y | 25280 | 24807 | 24575 | 23967 | 23843 |
| Sample093 | RedFox02 | Flank | Y | 24765 | 24172 | 23644 | 21972 | 21939 |

| | | | | | | | | |
|---|---|---|---|---|---|---|---|---|
| Sample094 | RedFox02 | Ext. Ear | Y | 16522 | 16042 | 15746 | 14815 | 14717 |
| Sample095 | RedFox02 | Groin | Y | 33869 | 31156 | 30881 | 28114 | 28064 |
| Sample096 | RedFox02 | Leg | Y | 25940 | 25339 | 25099 | 24173 | 24028 |
| Sample097 | RedFox03 | Axilla | Y | 22710 | 22373 | 22201 | 21812 | 21564 |
| Sample098 | RedFox03 | Flank | Y | 30133 | 29493 | 29324 | 27747 | 27414 |
| Sample099 | RedFox03 | Ext. Ear | Y | 21128 | 20669 | 20517 | 19110 | 19036 |
| Sample100 | RedFox03 | Groin | Y | 11925 | 11653 | 11480 | 11116 | 11050 |
| Sample101 | RedFox03 | Leg | Y | 26193 | 25011 | 24770 | 23553 | 23354 |
| Sample102 | RedFox04 | Axilla | N | 22057 | 21046 | 20877 | 20402 | 20399 |
| Sample103 | RedFox04 | Flank | N | 35549 | 34340 | 33876 | 31278 | 31180 |
| Sample104 | RedFox04 | Ext. Ear | N | 51241 | 49443 | 48641 | 45517 | 45256 |
| Sample105 | RedFox04 | Groin | N | 25295 | 24731 | 24493 | 23899 | 23705 |
| Sample106 | RedFox04 | Leg | N | 27010 | 26151 | 25980 | 25364 | 25223 |
| Sample107 | RedFox05 | Axilla | Y | 20807 | 20382 | 20306 | 20196 | 20160 |
| Sample108 | RedFox05 | Flank | Y | 10386 | 10161 | 9974 | 9587 | 9571 |
| Sample109 | RedFox05 | Ext. Ear | Y | 23427 | 23105 | 22946 | 22465 | 22411 |
| Sample110 | RedFox05 | Groin | Y | 9529 | 9245 | 9164 | 9005 | 9000 |
| Sample111 | RedFox05 | Leg | Y | 29618 | 29147 | 28964 | 27786 | 26951 |
| Sample112 | RedFox05 | Feces | Y | 37572 | 36118 | 35830 | 33293 | 32962 |
| Sample113 | RedFox06 | Axilla | N | 26804 | 26057 | 25638 | 24404 | 24247 |
| Sample114 | RedFox06 | Flank | N | 49845 | 48866 | 47369 | 42186 | 42049 |
| Sample115 | RedFox06 | Ext. Ear | N | 10965 | 10718 | 10393 | 9773 | 9755 |
| Sample116 | RedFox06 | Groin | N | 18970 | 18255 | 17677 | 13888 | 13772 |
| Sample117 | RedFox06 | Leg | N | 42852 | 41922 | 40979 | 36185 | 35033 |
| Sample118 | RedFox06 | Feces | N | 40038 | 38921 | 38633 | 36785 | 36575 |
| Sample119 | RedFox07 | Axilla | Y | 1810 | 1690 | 1569 | 1353 | 1353 |
| Sample120 | RedFox07 | Flank | Y | 1470 | 1383 | 1290 | 1137 | 1137 |
| Sample121 | RedFox07 | Ext. Ear | Y | 40520 | 39431 | 39242 | 38752 | 38483 |
| Sample122 | RedFox07 | Groin | Y | 65229 | 63691 | 63397 | 60378 | 58376 |
| Sample123 | RedFox07 | Leg | Y | 41419 | 40743 | 40597 | 40012 | 39611 |
| Sample124 | RedFox08 | Axilla | N | 26546 | 25817 | 25487 | 23929 | 23899 |
| Sample125 | RedFox08 | Flank | N | 57950 | 56300 | 55466 | 52325 | 51867 |
| Sample126 | RedFox08 | Ext. Ear | N | 30058 | 29223 | 28659 | 26911 | 26885 |

| | | | | | | | | |
|---|---|---|---|---|---|---|---|---|
| Sample127 | RedFox08 | Groin | N | 27783 | 26889 | 26303 | 24667 | 24562 |
| Sample128 | RedFox08 | Leg | N | 27233 | 26468 | 25858 | 23793 | 23582 |
| Sample129 | RedFox09 | Axilla | Y | 34948 | 34016 | 33849 | 33218 | 33141 |
| Sample130 | RedFox09 | Flank | Y | 30389 | 29933 | 29637 | 28443 | 28320 |
| Sample131 | RedFox09 | Ext. Ear | Y | 27316 | 26800 | 26601 | 26217 | 25778 |
| Sample132 | RedFox09 | Groin | Y | 37234 | 36322 | 36023 | 35606 | 35606 |
| Sample133 | RedFox09 | Leg | Y | 39676 | 38413 | 37968 | 35973 | 35574 |
| Sample134 | RedFox10 | Axilla | Y | 28930 | 28364 | 28022 | 27036 | 26925 |
| Sample135 | RedFox10 | Flank | Y | 33170 | 32197 | 31730 | 30387 | 30114 |
| Sample136 | RedFox10 | Ext. Ear | Y | 24691 | 23936 | 23781 | 23171 | 23171 |
| Sample137 | RedFox10 | Groin | Y | 29538 | 28969 | 28624 | 27860 | 27639 |
| Sample138 | RedFox10 | Leg | Y | 44650 | 43886 | 43524 | 42535 | 41992 |
| Sample139 | RedFox11 | Axilla | N | 6218 | 5989 | 5796 | 5215 | 5153 |
| Sample140 | RedFox11 | Flank | N | 34023 | 33074 | 32220 | 28765 | 28742 |
| Sample141 | RedFox11 | Ext. Ear | N | 13049 | 12657 | 12195 | 10853 | 10833 |
| Sample142 | RedFox11 | Groin | N | 23474 | 22555 | 22316 | 21190 | 21186 |
| Sample143 | RedFox11 | Leg | N | 42699 | 41395 | 41000 | 37965 | 37376 |
| Sample144 | RedFox12 | Axilla | Y | 31266 | 30772 | 30595 | 29882 | 29032 |
| Sample145 | RedFox12 | Flank | Y | 46195 | 44887 | 44448 | 40845 | 39906 |
| Sample146 | RedFox12 | Ext. Ear | Y | 48650 | 47339 | 46912 | 42652 | 42136 |
| Sample147 | RedFox12 | Groin | Y | 22500 | 21918 | 21737 | 21172 | 20700 |
| Sample148 | RedFox12 | Leg | Y | 46691 | 45758 | 45467 | 42561 | 42208 |
| Sample149 | RedFox13 | Axilla | N | 27153 | 26820 | 26294 | 24599 | 24217 |
| Sample150 | RedFox13 | Flank | N | 23406 | 23010 | 22466 | 20417 | 20206 |
| Sample151 | RedFox13 | Ext. Ear | N | 35123 | 34150 | 33217 | 29877 | 29660 |
| Sample152 | RedFox13 | Groin | N | 43264 | 42534 | 42132 | 40797 | 40047 |
| Sample153 | RedFox13 | Leg | N | 52624 | 51323 | 50902 | 48869 | 47900 |

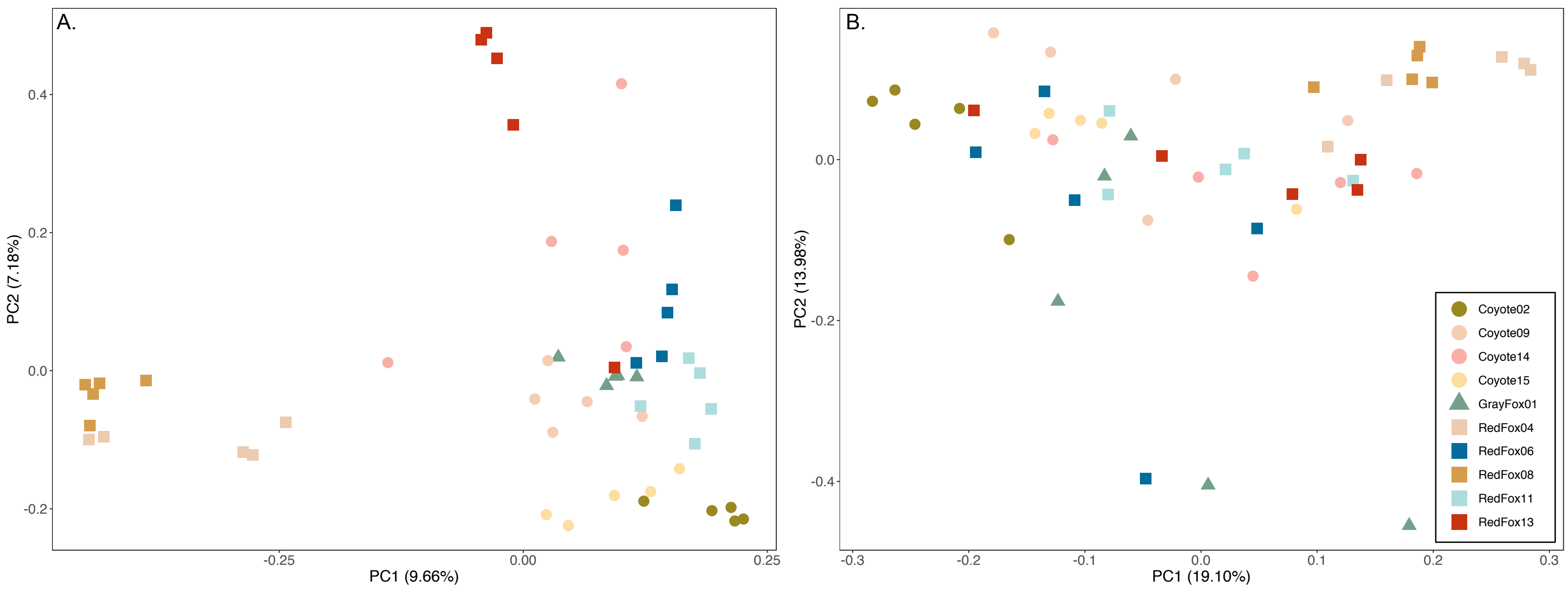


**Supplemental Figure S1.** Principal coordinate analysis (PCoA) of uninfected individuals showed significant clustering by individual (PERMANOVA; Bray-Curtis, *pseudo-F*= 2.984, *p*=0.001; Weighted UniFrac, *pseudo-F*=3.470, *p*=0.001) rather than body site (Bray-Curtis, *pseudo-F*=0.781, *p*=0.997; Weighted UniFrac, *pseudo-F*=0.950, *p*=0.574) using both (A) Bray-Curtis and (B) phylogeny-based Weighted UniFrac distances.

**Supplemental Table S2.** Results from alpha (Kruskal-Wallis test) and beta (PERMANOVA) diversity significance tests. Given that mange infection status exerted the strongest influence on alpha and beta diversity, we analyzed the full composite dataset in downstream analyses, with mange infection as our variable of interest.

**A.**

| | **Variable** | **Kruskal-Wallis *H*** | ***p*-value** |
|---|---|---|---|
| Chao1 | State | 4.879 | 0.300 |
| | Species | 2.468 | 0.291 |
| | Age | 2.771 | 0.428 |
| | Sex | 6.871 | 0.032 |
| | Year Sampled | 6.526 | 0.089 |
| | Mange | 10.711 | 0.001 |
| Evenness | State | 6.343 | 0.175 |
| | Species | 1.448 | 0.485 |
| | Age | 3.436 | 0.329 |
| | Sex | 6.546 | 0.038 |
| | Year Sampled | 4.659 | 0.199 |
| | Mange | 8.643 | 0.003 |

**B.**

| | **Variable** | ***Pseudo-F*** | ***p*-value** |
|---|---|---|---|
| Bray-Curtis | State | 1.341 | 0.015† |
| | Species | 1.025 | 0.358 |
| | Age | 1.075 | 0.271 |
| | Sex | 2.120 | 0.001 |
| | Year Sampled | 1.220 | 0.085 |
| | Mange | 3.885 | 0.001 |
| Weighted UniFrac | State | 1.324 | 0.135 |
| | Species | 1.244 | 0.22 |
| | Age | 0.880 | 0.627 |
| | Sex | 2.395 | 0.008 |
| | Year Sampled | 1.846 | 0.011† |
| | Mange | 4.398 | 0.001 |
| Unweighted UniFrac | State | 1.267 | 0.021† |
| | Species | 1.103 | 0.215 |
| | Age | 0.982 | 0.525 |
| | Sex | 1.554 | 0.007 |
| | Year Sampled | 1.230 | 0.047† |
| | Mange | 2.211 | 0.006 |

*† q-values from pairwise tests were not significant*

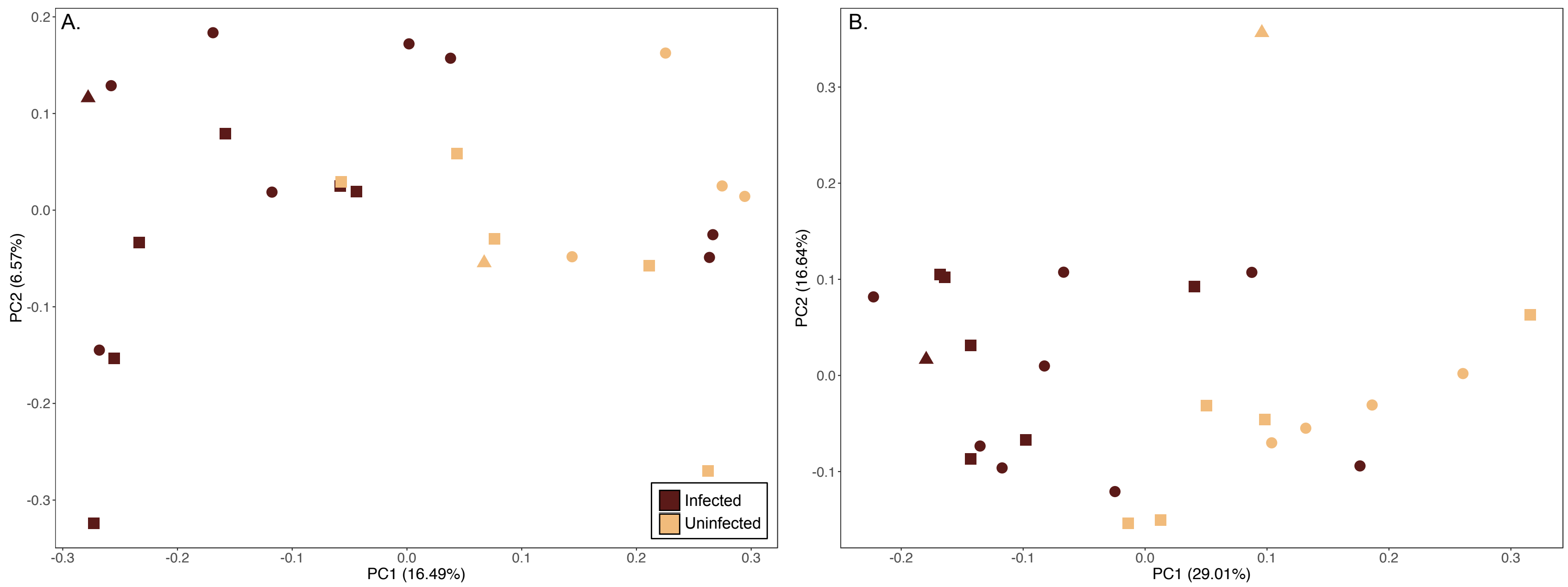


**Supplemental Figure S2.** Principal coordinate analysis (PCoA) showed significant differences between infection groups using both (A) Unweighted (PERMANOVA; *pseudo-F*=2.211, *p*=0.006) and (B) Weighted (*pseudo-F*=4.398, *p*=0.001) UniFrac distances.

**Supplemental Table S3.** Relative abundance of taxonomic groups by species and infection status. Across all three species, mange-infected individuals exhibited increased Bacilli and Actinobacteria, and decreased Other when compared to uninfected individuals.

| | Coyote | | Gray Fox | | Red Fox | |
|---|---|---|---|---|---|---|
| **Taxonomic Class** | **Infected** | **Uninfected** | **Infected** | **Uninfected** | **Infected** | **Uninfected** |
| Actinobacteria | 22.460% | 12.927% | 40.401% | 31.758% | 28.219% | 7.875% |
| Alphaproteobacteria | 3.642% | 9.956% | 0.673% | 5.291% | 0.831% | 3.530% |
| Bacilli | 32.249% | 5.523% | 26.874% | 0.747% | 42.096% | 14.602% |
| Bacteroidia | 4.147% | 2.974% | 2.621% | 0.145% | 1.367% | 2.649% |
| Betaproteobacteria | 1.196% | 2.608% | 0.380% | 0.403% | 1.277% | 2.972% |
| Clostridia | 6.806% | 4.487% | 12.417% | 0.240% | 2.291% | 17.902% |
| Flavobacteriia | 3.922% | 2.465% | 0.532% | 37.814% | 5.023% | 4.135% |
| Fusobacteriia | 4.111% | 4.340% | 5.303% | 4.291% | 0.717% | 3.969% |
| Gammaproteobacteria | 12.557% | 16.224% | 6.656% | 11.002% | 12.030% | 23.138% |
| Other | 8.911% | 38.496% | 4.143% | 8.310% | 6.148% | 19.229% |

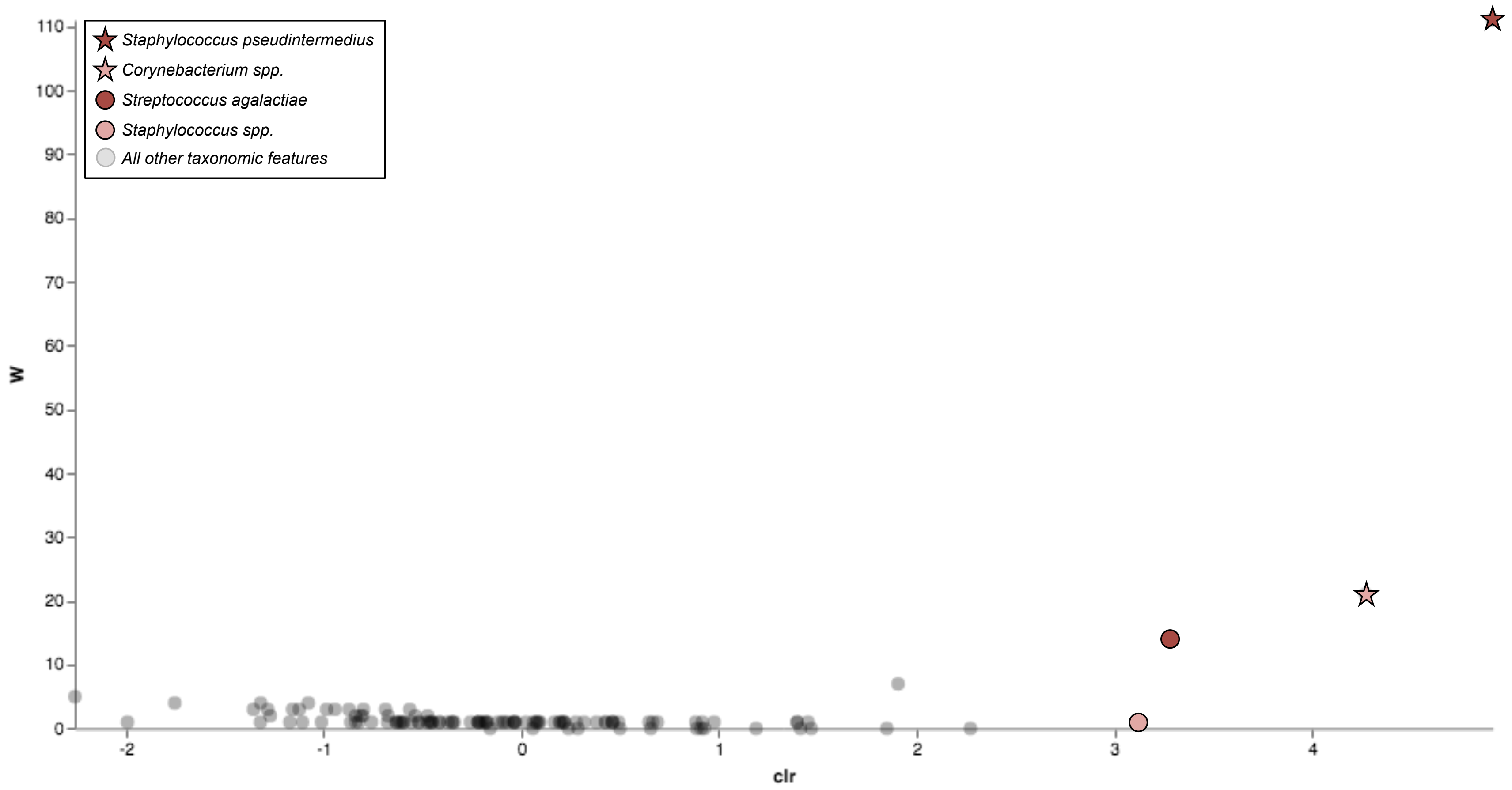


**Supplemental Figure S3.** Analysis of the composition of microbes (ANCOM) returned one taxonomic feature as consistently and significantly associated with mite infection status: *Staphylococcus pseudintermedius* (indicated with a red star). Three additional taxa that commonly co-occurred with *S. pseudintermedius* included *Corynebacterium spp.* (pink star), *Streptococcus agalactiae* (red circle), and *Staphylococcus spp.* (pink circle).

**Supplemental Table S4.** NCBI BLASTn results for the two features exhibiting increased relative abundance in mange-infected individuals: (A) 3f0449c545626dd14b585e9c7b2d16f4 (class: Bacilli) and (B) e3e89166daa575e51d7a14bc65f11153 (class: Actinobacteria).

| | **Species** | **Max Score** | ***E* value** | **% Identity** | **Accession No.** |
|---|---|---|---|---|---|
| **A.** | *Staphylococcus pseudintermedius* | 410 | 1.00E-110 | 100% | LC437049.1 |
| | *Staphylococcus pseudintermedius* | 410 | 1.00E-110 | 100% | LC437042.1 |
| | *Staphylococcus pseudintermedius* | 410 | 1.00E-110 | 100% | LC437038.1 |
| | *Staphylococcus pseudintermedius* | 410 | 1.00E-110 | 100% | LC437037.1 |
| | *Staphylococcus pseudintermedius* | 410 | 1.00E-110 | 100% | LC437027.1 |
| | *Staphylococcus spp.* | 410 | 1.00E-110 | 100% | MK954146.1 |
| | *Staphylococcus spp.* | 410 | 1.00E-110 | 100% | MK954144.1 |
| | *Staphylococcus pseudintermedius* | 410 | 1.00E-110 | 100% | MK681220.1 |
| | *Staphylococcus pseudintermedius* | 410 | 1.00E-110 | 100% | MK681219.1 |
| | *Staphylococcus pseudintermedius* | 410 | 1.00E-110 | 100% | CP035740.1 |
| | *Staphylococcus pseudintermedius* | 410 | 1.00E-110 | 100% | CP035741.1 |
| | *Staphylococcus pseudintermedius* | 410 | 1.00E-110 | 100% | CP035742.1 |
| | *Staphylococcus pseudintermedius* | 410 | 1.00E-110 | 100% | CP035743.1 |
| | *Staphylococcus pseudintermedius* | 410 | 1.00E-110 | 100% | CP032682.1 |
| | Uncultured *Staphylococcus spp.* | 410 | 1.00E-110 | 100% | MH728109.1 |
| **B.** | *Corynebacterium resistens* | 412 | 3.00E-111 | 100% | MF086686.1 |
| | Uncultured bacterium | 412 | 3.00E-111 | 100% | KX998044.1 |
| | Uncultured bacterium | 412 | 3.00E-111 | 100% | KF084913.1 |
| | Uncultured bacterium | 412 | 3.00E-111 | 100% | KF084799.1 |
| | Uncultured bacterium | 412 | 3.00E-111 | 100% | KF084787.1 |
| | Uncultured bacterium | 412 | 3.00E-111 | 100% | KF073670.1 |
| | Uncultured bacterium | 412 | 3.00E-111 | 100% | KF063189.1 |
| | *Corynebacterium resistens* | 412 | 3.00E-111 | 100% | NR_074826.1 |
| | Uncultured *Corynebacterium spp.* | 412 | 3.00E-111 | 100% | AB587824.1 |
| | Uncultured *Corynebacterium spp.* | 412 | 3.00E-111 | 100% | AB587823.1 |
| | Uncultured bacterium | 412 | 3.00E-111 | 100% | JX052081.1 |
| | Uncultured bacterium | 412 | 3.00E-111 | 100% | JX051886.1 |
| | *Corynebacterium auriscanis* | 412 | 3.00E-111 | 100% | EU911931.2 |
| | *Corynebacterium resistens* | 412 | 3.00E-111 | 100% | CP002857.1 |
| | Uncultured bacterium | 412 | 3.00E-111 | 100% | JF238803.1 |
